# The control of overt and covert attention across two nodes of the attention-control network

**DOI:** 10.1101/2024.01.05.574406

**Authors:** Pablo Polosecki, Sara C. Steenrod, Heiko Stemmann, Winrich A. Freiwald

## Abstract

Attention is a central cognitive capability whose focus is thought to be directed by a spatial map coding behavioral priority. Here we tested the three defining properties of priority map theory with electrophysiological recordings from two attentional control areas. Both areas, lateral intraparietal area LIP and dorsal posterior inferotemporal cortex PITd, selected behaviorally relevant locations even in the absence of visual stimuli (cognitive sustainment principle). Second, priority signals are thought to arise from the summation of multiple spatial signals (superposition principle). LIP approximated linear summation for visual stimuli, spatial attention, and eye movements, while PITd did not. Third, the same priority signal should guide different behaviors (agnosticity principle). LIP, instead, used separable processing channels for representing attentional focus and eye position, while PITd coded attentional focus only. Thus primate attentional control circuits implement multiple priority maps, whose functional diversity and dimensionality increase the computational capacity of attentional selection.

**SIGNIFICANCE STATEMENT:** A central theory of attention poses that the brain computes a priority map to highlight spatial locations of relevance for behavior. Here we tested the hypothesis and its key predictions about how priority signals are assembled and used, through electrophysiological single-unit recordings from two nodes of the attention control network, localized by functional magnetic resonance imaging. We found both areas to highlight locations even in the absence of a stimulus, and that each assembled spatial signals differently and provided spatial information in different forms. These findings force a revision of how and where spatial attention is controlled in the brain.

## INTRODUCTION

Areas of attention control in parietal and prefrontal cortex (1) contain spatial maps of the visual world (2–4). These maps are thought to encode a priority signal that can direct attention to a region of interest in space or a saccadic eye movement (5). The theory of priority maps includes a set of prescriptions for how priority is assembled from different visual and cognitive sources, and for the informational contents of the resulting signal in neural populations.

First, one basic property of attention control areas is their ability to highlight task-relevant spatial locations even in the absence of a visual stimulus (6–9), a feature we will refer to here as cognitive sustainment. This has frequently been shown as spatial tuning of neurons during the memory epoch in memory-guided saccade (MGS) tasks, where animals are required to make a saccade to remembered locations not signaled by a persistent visual stimulus. Cognitive sustainment has thus been proposed as characteristic of cognitive control, in contrast to earlier visual areas in which the attention gating is usually described as a function of the attention signal on the visual signal (10, 11). Cognitive sustainment has been reported across the set of brain regions thought to be part of the attention control network; in fact it has even been historically important for neurons to even be included in studies of cognitive control (12). Cognitive sustainment has not been tested, to the best of our knowledge, for inferotemporal areas, where recently a new attentional control area has been found (13).

Second, priority maps result from the integration of information from multiple channels (14). How are these input signals combined? Summation of overlapping spatial signals from visual regions and top-down signals related to goals and expectations has been proposed as a specific integration mechanism to generate priority signals (5). We refer to this as the superposition principle. It has originally been proposed for a region long thought to contain a priority map, the lateral intraparietal area (LIP) in the context of free-viewing visual search experiments (15). The superposition principle can also explain the interaction between spatial and feature-based attention (16). However, the decisive test of the superposition principle where cognitive spatial signals thought to be the core representation of priority maps (17) are independently controlled, has been lacking.

Third, the purpose of the construction of a priority map is that it would serve to support all kinds of goal-oriented behaviors. Priority signals are thus thought to be multi- purpose. They are also thought to be one-dimensional (18), lacking predetermined behavioral meaning (5). We refer to these properties as agnosticity (**Fig 1A**). Priority signals should be agnostic in the sense that they do not provide information about how they are used (19). Thus the same population activity generating a priority signal should be used to guide covert attention or overt attention (eye movements), whatever is appropriate at a given moment. The presumed lack of intrinsic information in population signals as to their eventual use has not been tested.

**Figure 1.**
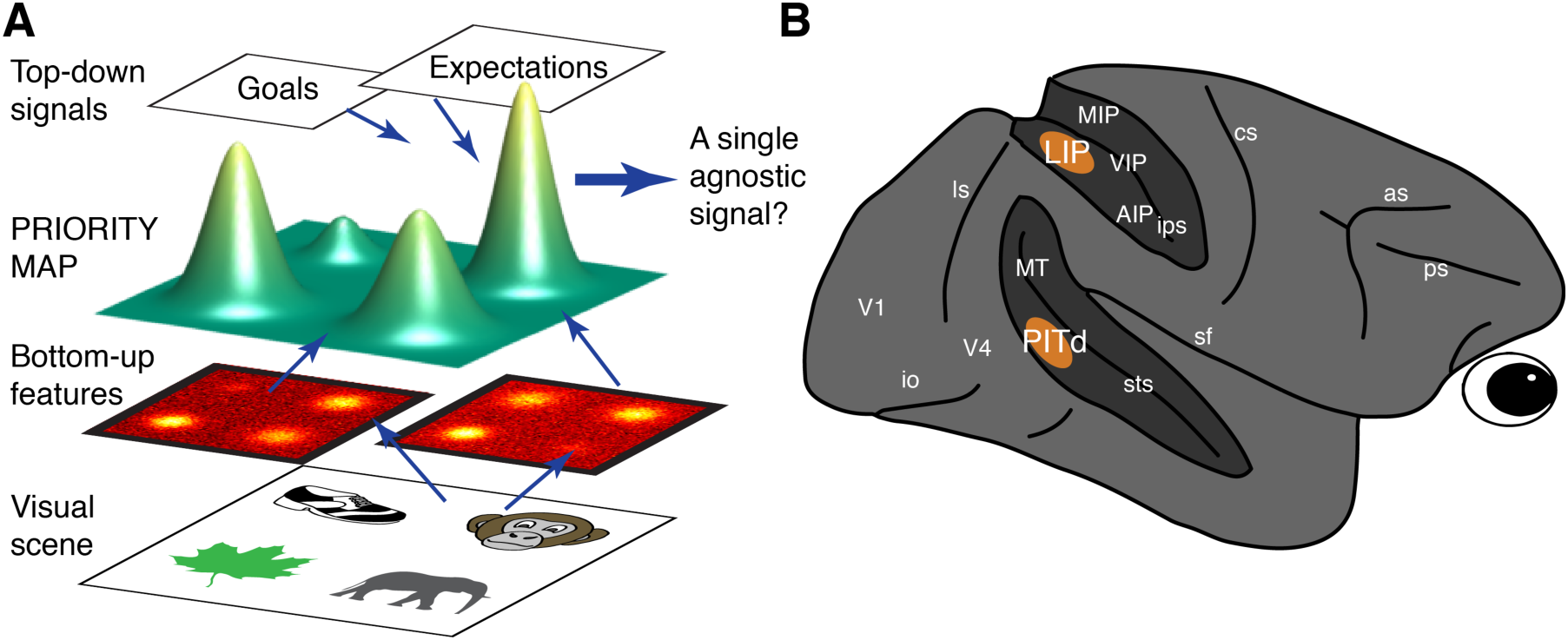
Conceptual framework and brain areas. (**A**) The priority map idea. A single spatial priority signal, agnostic as to its eventual use, is computed by summation of signals from feature-coding areas that represent visual-scene properties and internal signals related to goals and expectations. (**B**) Diagram of the macaque brain showing relevant sulci open and attention-control nodes localized by fMRI in areas PITd and LIP where recordings were performed.

These basic hypotheses, the cognitive sustenance hypothesis, the superposition and the agnosticity hypothesis, are central to the understanding of attention control. Yet they have not been directly tested together in a controlled experiment, nor directly compared across the different brain areas that are thought to implement them. Here we do this, taking advantage of the recent discovery of a new attention control area.

Recently, functional MRI (fMRI) has been used to map brain areas in the macaque brain modulated by attention (20). Beyond known attention control areas in parietal regions (LIP) and prefrontal regions (the frontal eye fields, FEF), the dorsal portion of posterior inferotemporal cortex (PITd) was strongly modulated by spatial attention, irrespective of visual features of relevant stimuli. Electrophysiological recordings confirmed this, and causal manipulations demonstrate that PITd activity controls the focus of spatial attention (13). This finding raises a fundamental question: Is this area in inferotemporal cortex really a priority map? The finding also offers a unique opportunity to determine the functional properties of priority maps and test the three core hypotheses about them. This is because PITd, unlike classical attention control areas like LIP (21) and FEF (4), has not been linked to eye movement control and might thus exhibit different functional properties from these classical attentional control areas.

How do the main principles of priority map theory compare in different locations of the attention control network? We tested the three principles of priority maps on areas PITd and LIP (**Fig. 1B**) using fMRI-guided extra-cellular recordings while monkeys performed a MGS task (for cognitive sustainment) and a novel behavioral paradigm that dissociates covert attention and eye movements (for superposition and agnosticity).

## RESULTS

### fMRI-guided electrophysiology

We performed extracellular recordings in areas PITd and LIP from two male rhesus monkeys (*macaca mulatta*, M1 and M2). Recording locations were chosen based on fMRI maps of spatial attention effects (20) in the same animals (Fig S1). We recorded from 85 PITd and from 66 LIP units, respectively. The only selection requirement for cells was that their receptive fields (RFs) had to be non-foveal (see Methods).

### Cognitive sustainment

We first measured the ability of each area to highlight task-relevant spatial locations in the absence of visual stimuli during a standard MGS task (See Methods, **Fig. 2A**). We computed spike density functions (SDFs) in single units and found units with spatially tuned activity throughout the duration of the trial (**Fig. 2B**). Large fractions of LIP and PITd cells, 73% and 40%, respectively, were significantly tuned during the memory epoch of the trial (600 to 400ms before saccade onset, **Fig 2C**). Selectivity was significant in a larger fraction of the recorded LIP units than of PITd units, and the range of effects strengths was larger in LIP than PITd. Yet both areas reflected pure spatial selection. The similarity of responses in both areas is surprising given their very different locations within the visual system (22), one part of the dorsal, the other of the ventral stream. Results from PITd suggest strictly cognitive signals, reflecting covert attention to a location with no stimulus, exist in an inferotemporal cortical area. This property, up to now a hallmark of higher cognitive and oculomotor areas, is expected from a priority map.

**Figure 2.**
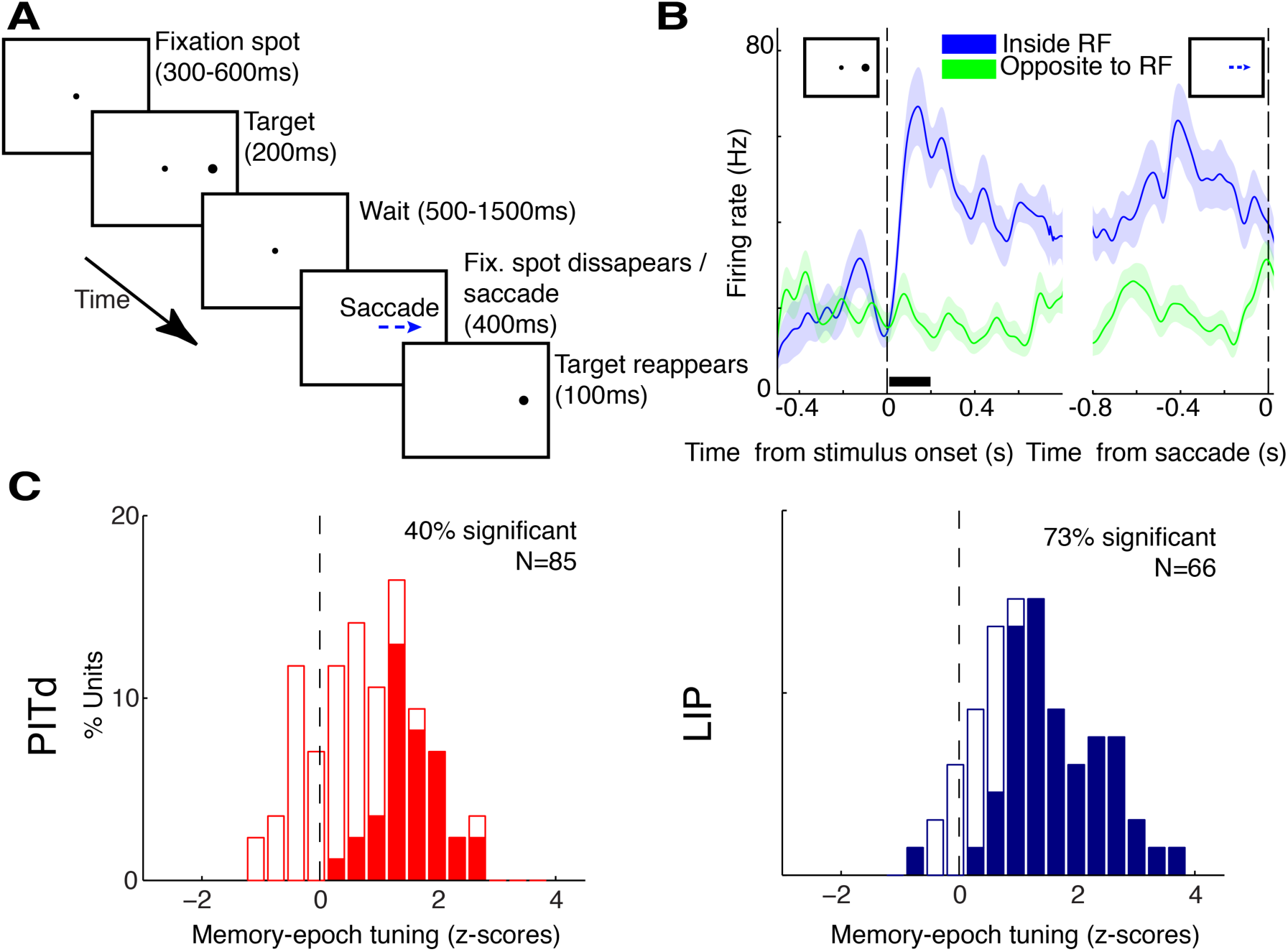
Cognitive sustainment: memory-guided saccade (MGS) task and neural responses. (**A**) Timing of events in the MGS task. See main text and Supplemental Methods for task description. (**B**) Spike density function (SDF) of an example PITd unit, aligned to stimulus or saccade onset. Color indicates stimulus position relative to the receptive field (RF). Shading denotes standard error of the mean (SEM). Horizontal bar indicates stimulus duration. (**C**) Distribution of strength of spatial tuning during the memory epoch (600 to 400ms before saccade onset) in PITd (left) and LIP (right). Filled bars indicate units with statistically significant tuning (t-test) at a false discovery rate q<0.05 in each area. Units from both monkeys were pooled together.

### Behavior during a covert motion discrimination task

For the study of superposition and agnosticity, we used a second task. The defining feature of this covert motion discrimination (CMD) task (**Figs. 3A, B**) was that the loci of covert and overt spatial attention were independently manipulated. In brief, subjects were required to attend one of two moving-dot surfaces (MDSs), while keeping fixation on the central fixation point (FP), in order to be able to report the motion direction of the cued MDS (the target stimulus) with a saccade to a saccade target dot (ST). Trials started with a central fixation period after which a small central cue indicated the location of the task-relevant MDS appearing 2s later (**Fig. 3A**). In each MDS, only a fraction of dots moved coherently in the same direction, and the direction of motion changed every 60ms (independently for each MDS) for a variable period of time (60 to 3600ms, also independent for each MDS) until the occurrence of the task-relevant event. These features required subjects to pay sustained covert attention over long periods of time, without any information about the direction of the saccade they would eventually need to generate. The task-relevant event was a prolonged motion event (PME) with constant motion direction, which lasted up to 2.7 seconds. (PME timing and direction were independent for each MDS.) PME motion directions were in one of two opposite directions, and monkeys had to indicate the motion direction in the target MDS, as soon as they made a decision, with a saccade to a ST. Monkeys successfully performed this task and, specifically, responded to the PME in the cued MDS, while ignoring that in the distractor (**Figs. 3C, 3D, S2** and Supplemental Information “Behavior Results”).

**Figure 3.**
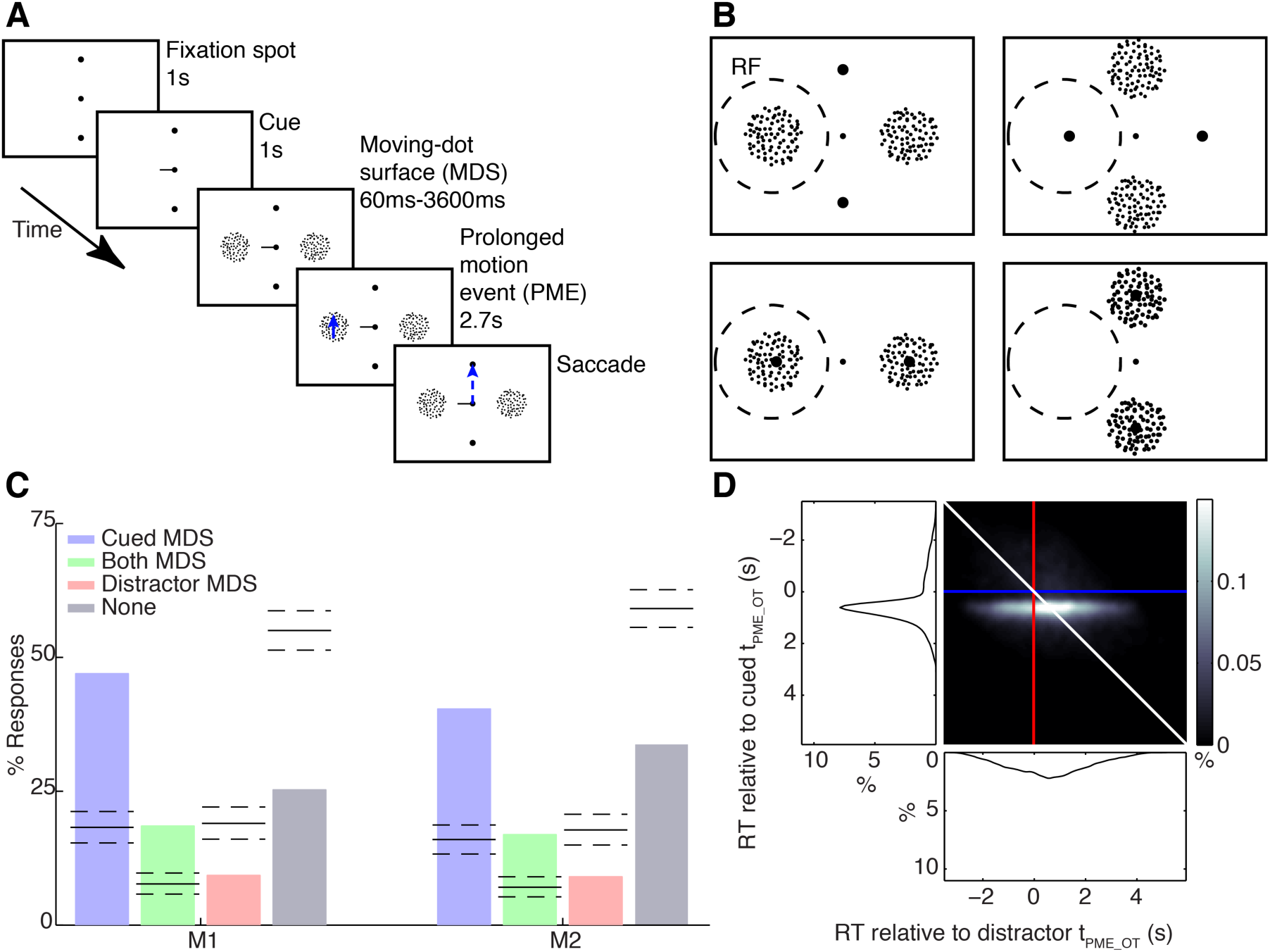
Covert discrimination task and behavioral performance. (**A**) Timing of events in the covert-discrimination task. (**B**) The four physical configurations of the stimuli relative to the RF of the unit being recorded: a lone moving-dot surface (MDS), a lone saccade target, both overlapping, or none. See main text and Supplemental Methods for task description. (**C**) Consistency of behavioral responses in monkeys M1 and M2 to timing and direction of the PME in the cued MDS only, both MDSs, distractor MDS only, and none of them. The first two cases correspond to correct trials. Lines indicate average chance levels and 95% confidence intervals (randomization test) across experimental sessions. (**D**) Joint distribution of response times (RTs) of monkey M1 relative to the prolonged motion event (PME) onset time (tPME_OT) in the cued and distractor MDS. See also Figure S2.

Crucially for the purposes of this study, the spatial position of task-relevant elements (MDSs and STs) relative to the RF of the recorded neuron, was changed across trials into four unique physical stimulus arrangements (**Fig. 3B**): an MDS as the only stimulus inside the RF, a ST as the only stimulus inside the RF, both an MDS and a ST overlapping inside the RF, and neither an MDS nor a ST positioned inside the RF. For each spatial arrangement, there were four unique behavioral situations (two possible cued MDSs and two possible saccade directions), making a total of 16 unique conditions overall, where all spatial combinations of covert and overt cognitive spatial signals were exhaustively sampled.

### Combination of spatial signals

Neurons obeying the superposition principle summate incoming signals. This property can be tested by determining the responses of a neuron to conditions with a single spatial signal inside their RF (response rA when covert attention is paid into its RF, response rS when a saccade is generated into its RF) and the response to conditions when both signals overlap in the RF. For a neuron to obey the superposition principle, this response should be the sum of rA and rS. An example for how such a neuron should behave in the CMD task is shown in **Fig 4A** (top row). Neurons in both areas exhibited a large response repertoire including modulation by covert attention, pre-saccadic tuning, and their combination (**Figs. 4A**, second row, **and S3** for examples). Population-average SDFs (**Fig. 4A**, third and bottom row for PITd and LIP, respectively) demonstrate widespread attention and saccadic modulations in both areas. Thus PITd, like LIP, is modulated by overt attention.

**Figure 4.**
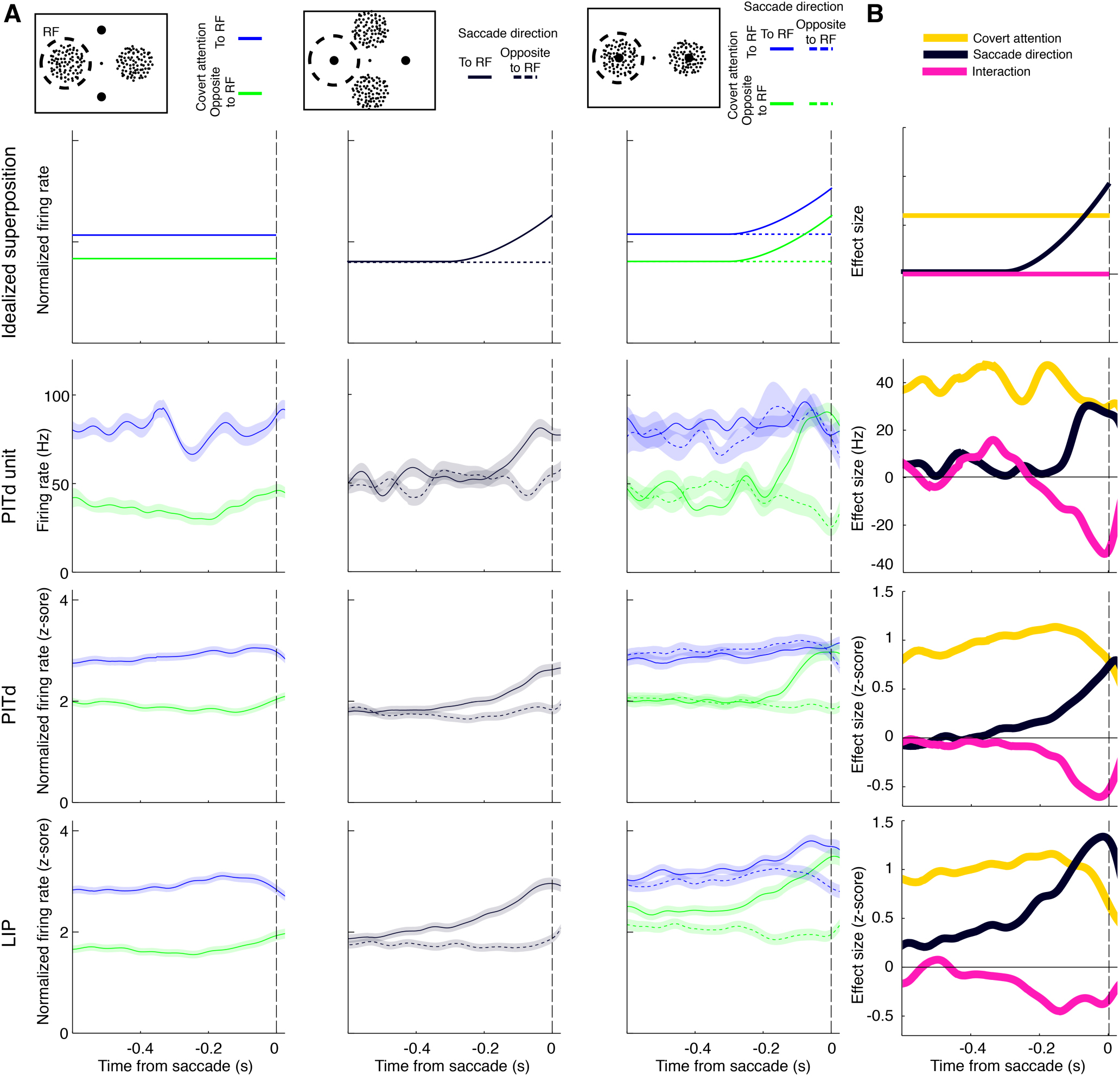
Testing superposition of spatial signals in PITd and LIP. (**A**) Top row: Idealized schematics of the superposition principle of neural responses on the covert discrimination task, showing a SDF, with activity aligned to saccade onset. Signals from covertly attending a lone cued MDS in the RF (left) and planning a saccade to a lone target (center) combine additively when these overlap in space (right), with both attention and saccadic responses observed. Second row: SDF of an example PITd unit with non-additive responses. Shadows indicate standard error of the mean (SEM). Third row: Population average of normalized (z-score units) PITd SDFs. Bottom row: Population average for LIP. (**B**) Main effect of covert attention, of eye movement conditions, and their interaction, in a generalized linear model (GLM). Top row: Idealized superposition showing no interaction between spatial signals. Second row: same PITd unit as in (**A**). Third row: PITd population average. Bottom row: LIP population average.

For the superposition principle to hold, saccade and attention signals should be independent of each other. Surprisingly though, they were not. In particular, saccade signals depended strongly on the attention condition. This deviation was not just a quantitative deviation from exact summation, but qualitative: in particular, no saccadic signals existed in PITd when the attended MDS was inside the RF. In contrast, they still existed in LIP, although reduced (**Fig. 4A**, third and bottom row).

To characterize the contributions of physical (presence of stimuli in the RF) and cognitive factors (attention and pre-saccadic tuning), we computed a time-resolved generalized linear model (GLM) of neural activity temporally aligned to saccade onset. Interaction terms in the GLM directly quantified the size of nonlinearities or deviations from superposition (which predicts no interactions between spatial signals, **Fig. 4B**, top). We calculated time courses of these effects in individual units and their normalized population average (**Fig. 4B**, **Fig. S4A**). Both areas exhibited strong effects of covert attention (PITd, mean±SEM in z-score units: 0.85±0.06, LIP: 0.88±0.09; see Methods) and saccade tuning (PITd: 0.40±0.06, LIP: 1.0±0.1) as well as a nonlinear interaction between them (PITd: -0.39±0.06, LIP: -0.37±0.06). The interaction was negative, indicating signals on average combined sub-additively. Its size in PITd was so big, it was of the order of main saccade tuning effect. In LIP, interaction effects were small compared to saccade tuning and covert attention. We next determined strength and significance of these effects for each neuron separately, and then computed the distribution of all effects across the population in each area (**Fig. S4B**). Large fractions of neurons were significantly affected by each task variable. Interactions between covert attention and saccade direction were common in PITd (significant in 21% of neurons) and less common and significant in LIP (0% of neurons). These results provide strong evidence for a systematic interaction between two kinds of spatial cognitive signals in PITd and thus against the superposition principle operating in this attention control area. The results also provide strong evidence that spatial cognitive signals are superposed, although attenuated, in area LIP.

### Spatial information represented by neural populations

According to the agnosticity principle, priority maps should provide no intrinsic information about the behavioral use of the combined spatial signals in the RF. The independent control of spatial orienting behaviors in the CMD task enabled us to test this prediction by measuring the simultaneous information about them in neural populations in PITd and LIP. We quantified the presence of information by the decoding accuracy of support vector machines using single trial activity of pseudopopulations (23) aligned to saccade onset (see Methods). We attempted to decode the covertly attended location when an MDS was alone in the RF, and the saccade direction when a ST was alone inside the RF. Crucially, we repeated the analyses when the MDS and ST overlapped in the RF. If agnostic, the resulting spatial signals should simply indicate the behaviorally relevant location at each time: the covertly attended location initially, then the upcoming saccade (**Fig 5A**, left).

**Figure 5.**
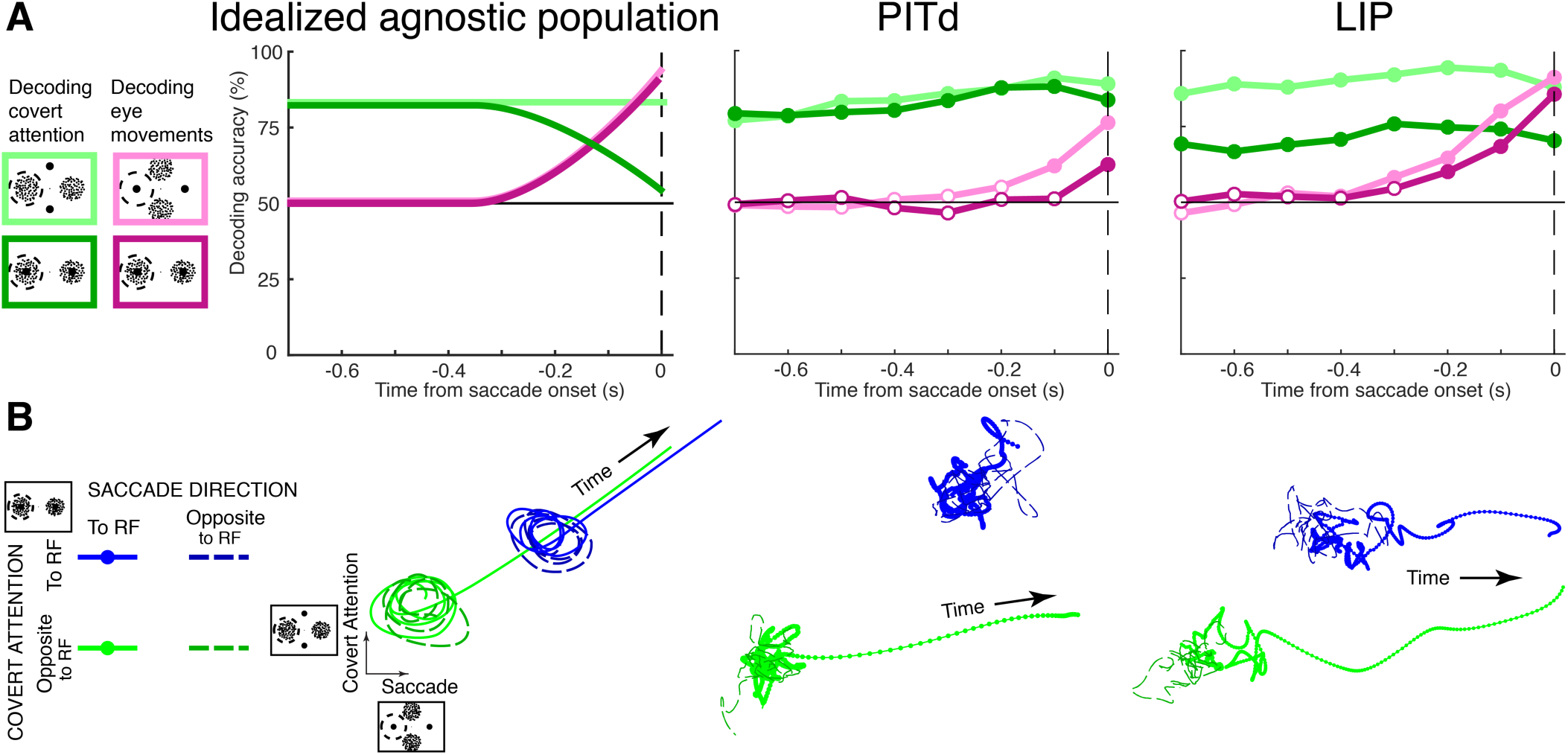
Testing agnosticity of population activity in PITd and LIP by decoding cognitive variables in the covert discrimination task. (**A**) Left: Accuracy, aligned on saccade onset, expected on an idealized population with an agnostic priority signal for decoding covert attention location (green) and saccade direction (pink) when single stimuli are inside the RF (light trace) or both are (dark trace). Center: Decoding accuracy for PITd, in 100ms time bins. Right: Decoding accuracy for LIP. Circles indicate the end of a time bin. Filled circles denote performance significantly different from chance levels (p<0.05, permutation test, Holm-Bonferroni controlled for comparisons on multiple time points). (**B**) Population activity when both stimuli are inside the RF of cells, projected on axes associated with covert attention conditions (vertical) and saccade direction (horizontal). Color denotes covert attention condition; and line style, saccade direction. Left: Idealized one-dimensional priority signal. See Fig. S5 for other alternative scenarios. Center: PITd population. Units (z-scores, not shown) are the same for vertical and horizontal axes. Right: LIP population.

In PITd, when single stimuli where in the RF, attention could be decoded significantly better than chance at all times (**Fig. 5A, center**, light green), and saccadic information at the end of the trial (**Fig 5A, center**, pink). In the overlapping condition, decoding of covert attention remained practically unchanged (dark green), and decoding accuracy of saccade direction was weaker (purple). Thus, the observed interactions in PITd (**Fig. 4B**, **Fig. S4A**) come at the expense of information loss about saccades. The resulting signal in PITd is not agnostic, but robustly signals the locus of covert attention. In LIP, covert attention location could be decoded at all times for both stimulus arrangements (**Fig. 5A, right**, light and dark green). Notably though, decoding of covert attention was degraded by the presence of a saccade target inside the RF, and it was degraded at all times (dark versus light green lines), even before any information about the saccade target could be decoded. This information about saccade direction increased until saccade onset reaching similar and significant levels for both stimulus arrangements (pink and purple lines). Thus LIP accommodates information about attention and saccades simultaneously, sacrificing attentional signal-to-noise. Crucially, these results rule out the agnosticity principle in both areas.

LIP representations have been suggested to be one-dimensional (18). How then is spatial information about covert and overt attention (**Fig. 5A**) accommodated in the population? To answer this question, we used targeted dimensionality reduction (24) to visualize dynamics in population space (see Methods). For each population (PITd and LIP), GLM regression coefficients from trials with MDSs *alone* inside the neurons’ RF were used to compute a population direction associated with covert attention. Analogously, regression coefficients from trials with saccade targets *alone* in the RF were used to isolate a direction associated with saccade direction. Into the plane these two axes span, we then projected the mean time course in trials when *both* MDSs and STs were present inside the RF (**Figs. 5B**). An agnostic representation resulting from superposition would be one dimensional (**Fig. 5B, left**). Motor representations would also be one-dimensional, but show different dynamics (**Fig. S5**). PITd population activity deviated from this scenario in multiple ways (**Fig 5B, center**). First, PITd population activity showed no saccadic contribution when attention was inside the RF. Second, activity during the two covert attention conditions was separable at all times. In LIP attention and saccadic spatial signals evolved along each population axis independently (**Fig 5B, right**). Thus LIP population activity was not one-dimensional, but exhibited separable dynamics for overt and covert attention.

These results clarify the nature of cognitive signal representations in two attentional control areas. In PITd, covert attention signals dominate. Saccade directions, in contrast to attention signals, are not reliably represented. In fact, the PITd population lacks a representation of saccade-direction when attention is already directed to the RF. A purely attentional representation (**Fig. S5**), comes closest to PITd population dynamics. In LIP, the meaning of axes computed from single-stimulus conditions remained valid when spatial signals overlapped in space (**Fig 5B, right**). This shows how the superposition principle can apply to a population code, while the resulting priority signal is entirely “gnostic”, allowing to tell overt and covert attention apart. Thus, we find empirically that the superposition principle preserves rather than dilutes the behavioral meaning of each signal, close to what is expected in a full dissociation scenario (**Fig. S5**).

## DISCUSSION

We have tested three fundamental principles of priority maps in two fMRI-identified attention control areas: one, PITd, whose role in attention control was recently discovered (13, 20) and another, LIP, for which many of the priority map ideas had been originally proposed but not systematically tested. The demonstration of cognitive sustainment in PITd puts it in a comparable position with established attention control nodes such as LIP. Yet, differences in superposition and information content suggest diverse functional roles of the contributions of both areas to the control of spatial attention. Strong nonlinearities in PITd invalidated the superposition principle and instead provide a robust readout of covert attention. In LIP, the superposition principle holds, and rather than providing an agnostic one-dimensional signal it allows simultaneous representations of cognitive spatial information to coexist.

### Cognitive Sustainment

Cognitive sustainment of neural activity is a defining characteristic of parietal (6), prefrontal (7, 8), and subcortical attention (9) control nodes. These areas are all part of the oculomotor system, where cognitive sustainment has been used as a criterion for identifying cells not involved in exclusively in motor planning (12, 25). The finding of the same signals outside of this system, in inferotemporal cortical area PITd, shows that cognitive sustainment can exist outside of the oculomotor system and is thus not tied to saccade planning. The finding, furthermore, differentiates area PITd from the standard view on early and ventral stream areas as recipients of attentional gating signals modulating visual activity (11, 26). Instead the results support the new hypothesis (13) that PITd is a possible source of cognitive spatial signals in the temporal lobe, yet one not involved in saccade planning.

The finding of cognitive sustainment in PITd shows how, in many ways, this temporal lobe area is functionally similar to parietal area LIP. Below we are discussing functional differences between the areas, but the results demonstrate how both are coding for visual stimuli, for covert, and for overt attention. The finding of an area located right next to two face-selective areas (20), to share properties with other attention control areas is surprising. It suggests that LIP and PITd might be directly connected to each other to negotiate a joint focus of attention and how PITd might serve to relay priority signals it has computed into other attentional control areas that are part of the oculomotor system. Our results thus force a re-conceptualization of attention control networks. And they force a network-oriented thinking where the major functions of an area can be highly different from those of neighboring areas, yet similar to more distant areas in the brain.

### Superposition of spatial cognitive signals

Linear superposition has been proposed as a mechanism for the formation of a spatial priority signal (5, 15). To our knowledge, this is the first time this hypothesis has been tested for spatial cognitive signals, arguably the core computation of priority maps (17), in a experiment that manipulated them independently, in any node of the attention network. In LIP, we find spatial signals can be attenuated when overlapping, yet their presence is preserved, largely validating the superposition principle. This offers a general prescription for how activity observed in simple experiments (as trials with a single stimulus in the RF can be considered) can predict that in more elaborate scenarios, perhaps indicating an effective limit to the complexity they could reveal (27). In PITd however, superposition fails. This indicates that there is not one mechanism for the formation of attentional control areas, and suggests how each area might read out signals originating elsewhere in the network while preserving its functional identity.

### Agnosticity of population representations

The idea of an agnostic general signal has been suggested as a key property of priority maps (19, 28). It has also been proposed to result directly from superposition (15): if multiple sources are combined in neuron responses, there might not be a predefined behavioral meaning to them. Results in both PITd and LIP rule out a general agnostic signal. In PITd only information about the attended location was robust. This confirms expectations for an area outside of the oculomotor system, and instead suggests a role as a true spotlight of attention (**Fig. 6, left**) for adjacent-feature coding areas, consistent with causal interventions (13). The lack of agnosticity in LIP is surprising, since we observed the expected superposition of spatial signals there. Key for understanding this is that even if superposition were to take place perfectly, there is heterogeneity in neural responses (12). This differential selectivity can preserve information in ways that are simple but not obvious in responses of individual neurons. Superposition, then, a straightforward encoding scheme for neuron responses, preserves the nature of contributing signals. Because of it, multiple parallel channels of spatial information coexist (**Fig. 6, right**), perhaps allowing LIP to function as a versatile look-up table for the variety of brain regions to which it is connected and the behaviors they support.

**Figure 6.**
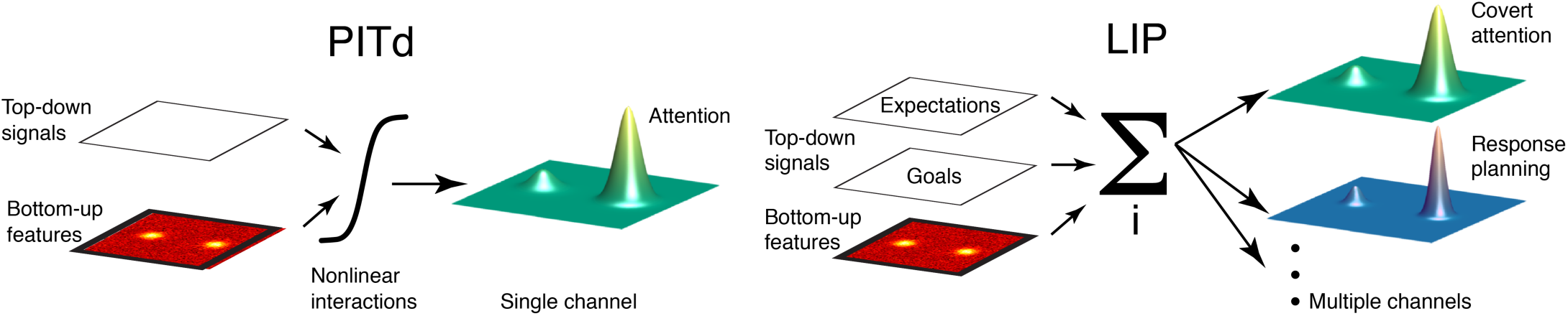
Schematics of functional specialization of attention-control nodes. Left: Coding of attention in PITd, where nonlinear interactions dominate. Right: Simultaneous representation of multiple spatial variables in LIP at the population level through sum of disparate signals.

The present findings also bear on the recently growing interest in understanding the dimensionality of neural representations (23, 24). The study of attention and decision-making in separate experiments has led to the proposal that responses in LIP evolve along a one-dimensional manifold (18) but it has been recently argued that additional dimensions can only be revealed in more complex tasks (27). In that sense, the superposition principle provides a linear embedding of task parameters in neural space, in contrast with higher cortical areas where interactions account for the richness of the representation (23) needed for learning arbitrary associations.

Taken together, the results highlight the role of PITd as an attention control area in the temporal lobe with a relatively specialized representation for attention robust to perturbations from other cognitive sources, and that of LIP as holding as a multidimensional map able to guide perception and action at once.

## METHODS

Two adult male rhesus monkeys (10 and 8 kg) were trained on a memory-guided saccade (MGS) and a covert-discrimination tasks. All animal procedures complied with the US National Institutes of Health *Guide for Care and Use of Laboratory Animals* and were approved by The Rockefeller University Institutional Animal Care and Use Committee (IACUC). While the monkeys were engaged in the behavioral tasks, we recorded single- and multiunit responses in temporal area PITd and parietal area LIP, targeting attention-sensitive hotspots measured with functional MRI in a previous study on the same animals (20). The great majority of neurons were not recorded simultaneously, but rather in separate behavioral sessions. For the MGS task, we quantified the existence of spatial tuning during the memory period. For the covert- discrimination task, we first analyzed the behavioral responses to verify monkeys were reliably using the cue for covertly guiding their responses and ignoring the distractor stimulus, and we then analyzed neural responses. First, we quantified the effect in each unit of the spatial signals manipulated during the task as well as their interaction. We then pooled data from single- and multiunit recordings to construct population responses, and used support vector machines to decode different covert attention and direction of impending saccades. We applied a dimensionality reduction technique (‘targeted dimensionality reduction’) to identify a low-dimensional subspace capturing variance due to the task variables of interest within population responses. Full methods are provided in the Supplemental Information.

## ACKNOWLEDGMENTS

We thank P. Levy and M. Borisov for help with data analysis; R. Huq for help with experimental planning; L. Diaz, A. Gonzalez, S. Rasmussen, and animal husbandry staff of The Rockefeller University for veterinary and technical care. This work was supported by the National Institute Of Mental Health of the National Institutes of Health under Award Number R01MH120288 and the National Science Foundation under Award Number BCS-1057006. W.A.F. is a NYSCF-Robertson Investigator. P.P. received financial support from the David Rockefeller Graduate Program. The content is solely the responsibility of the authors and does not necessarily represent the official views of the National Institutes of Health and the National Science Foundation.

## AUTHOR CONTRIBUTIONS

P.P., S.C.S., and W.A.F. conceived experiments. H.S. provided experimental resources. P.P. and S.C.S. performed experiments. P.P. analyzed experiments. P.P. and W.A.F. wrote the manuscript.

## Supplemental Information

### SUPPLEMENTAL FIGURES

**Fig. S1.**
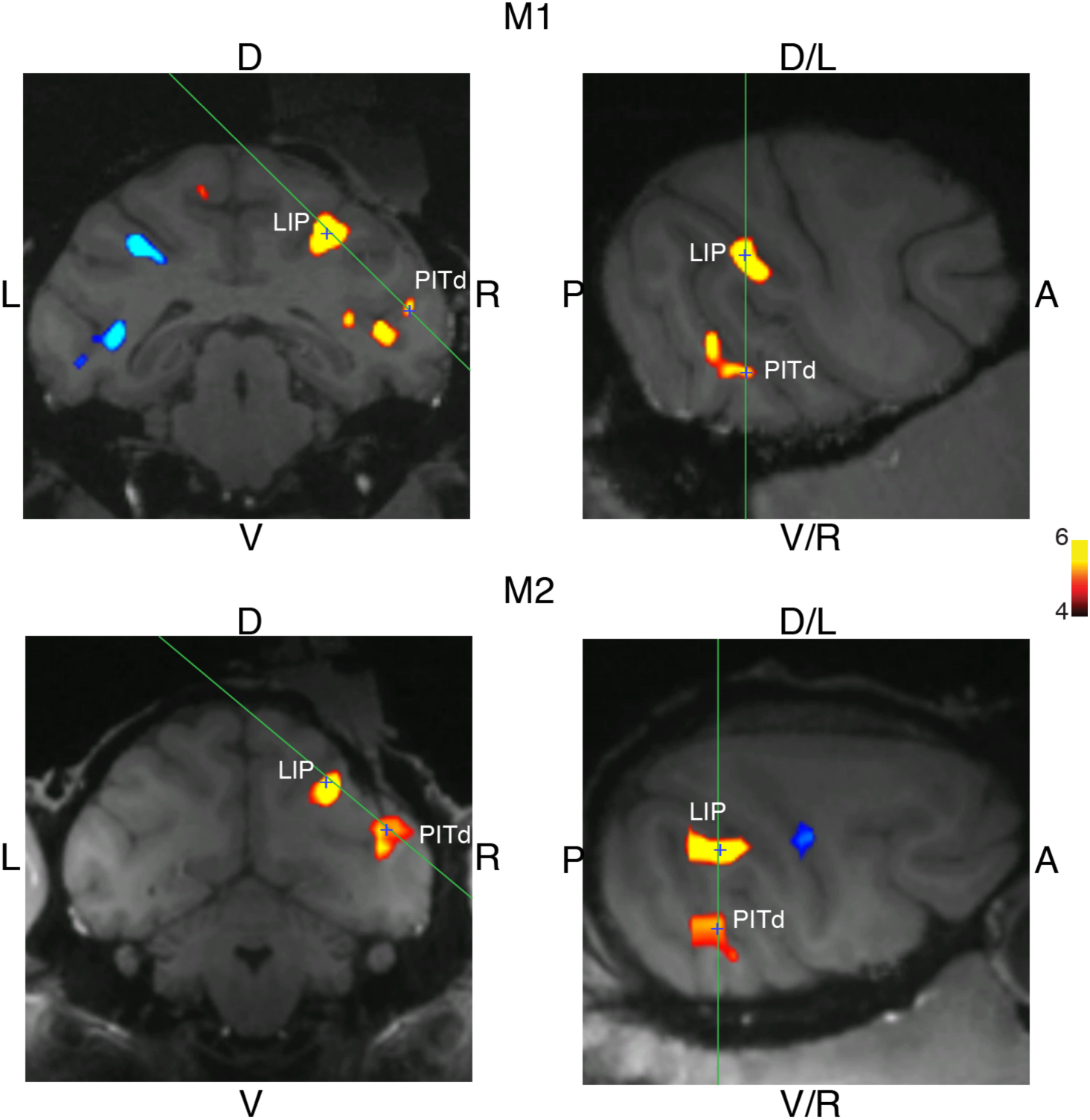
fMRI-guided electrophysiology. Coronal and sagitto-horizontal planes showing recoding locations (marked with a blue cross) targeting PITd and LIP in monkeys M1 and M2. Green lines indicate the orientation of the related slice in the adjacent panel. Colored overlay is a thresholded statistical parametric map of covert attention effects (20) used to guide recordings (t-statistic, scale bar on right side). PITd location in monkey M1 is 0mm anterior to the interaural line (+0, AP) axis, +23 to the right on the medio-lateral (ML) axis and +18 dorsally along the dorso-vental (DV) axis. LIP in monkey M1 was targeted at AP +0, ML +12, DV +28. PITd in monkey M2 was targeted at AP -1.5, ML +23, DV +21. LIP in monkey M2 was targeted at AP - 2, ML +14, DV +29.

**Fig. S2.**
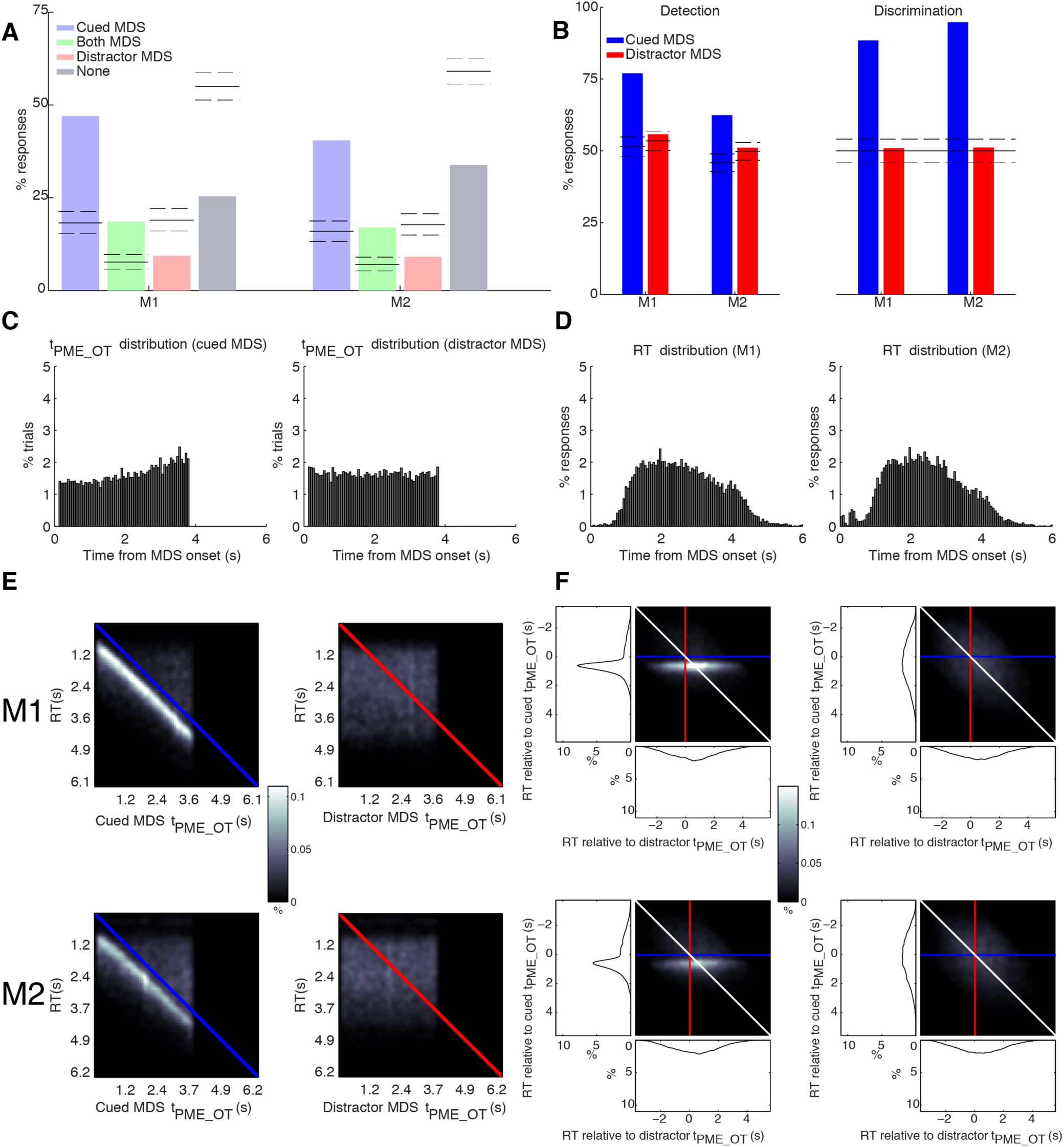
Analysis of behavioral performance in monkeys M1 and M2. (Corresponds to Figure 3) (**A**) [Identical to Fig. 3C] Consistency of behavioral responses in monkeys to timing and direction of the PME in the cued MDS only, both MDSs, distractor MDS only, and none of them. The first two cases correspond to correct trials. Horizontal lines indicate average chance levels and 95% confidence intervals (randomization test) across experimental sessions. (**B**) Left: Average detection performance across experimental sessions with respect to cued MDS (blue) and distractor MDS (red). Right: Average discrimination performance across experimental sessions. (**C**) Frequency distribution of PME onset times (tPME_OT) relative to MDS onset. Left: cued MDS. Right: distractor MDS. (**D**) Frequency distribution of response times (RTs) Left: monkey M1. Right: monkey M2. (**E**) Joint distribution of RTs and tPME_OT of the cued (left) or distractor (right) MDS in monkeys M1 (top) and M2 (bottom). (**F**) Left: Joint distribution of RTs measured relative to tPME_OT values of cued (y axis) and distractor (x axis) MDSs in the same trials. Histograms showing marginal distributions are shown adjacently. Right: Similar to left panel, but using shuffled-trial RT values instead of same-trial RTs (null distribution). The distribution shown is the average across 10^5^ reshuffles.

**Fig. S3.**
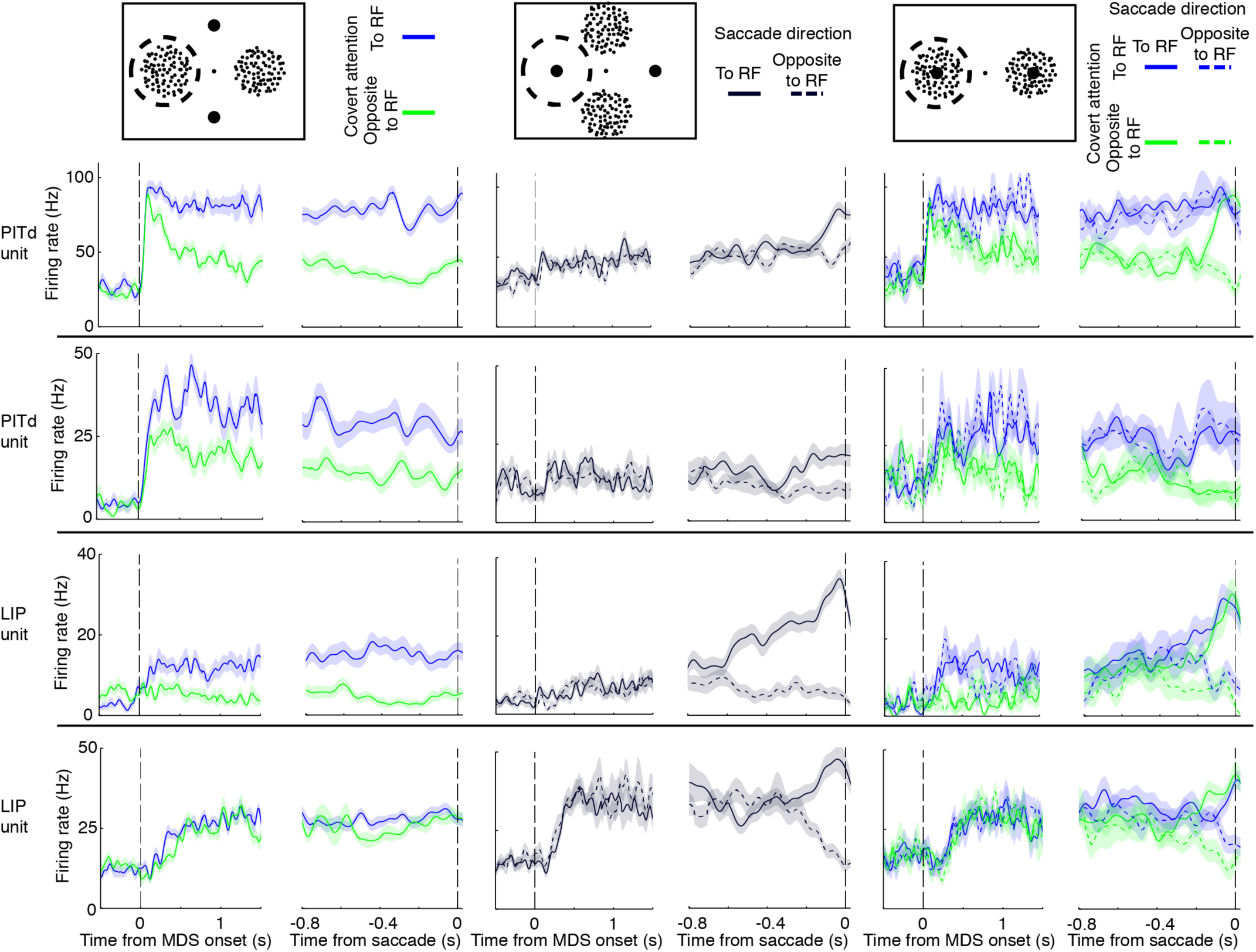
Neural responses during the covert-discrimination task. (Corresponds to Figure 4A) SDFs of example units (top row is the same unit as Fig. 4A) for areas PITd and LIP, time aligned on MDS onset (left half) and saccade onset (right half). Conventions as in Fig. 4A.

**Fig. S4.**
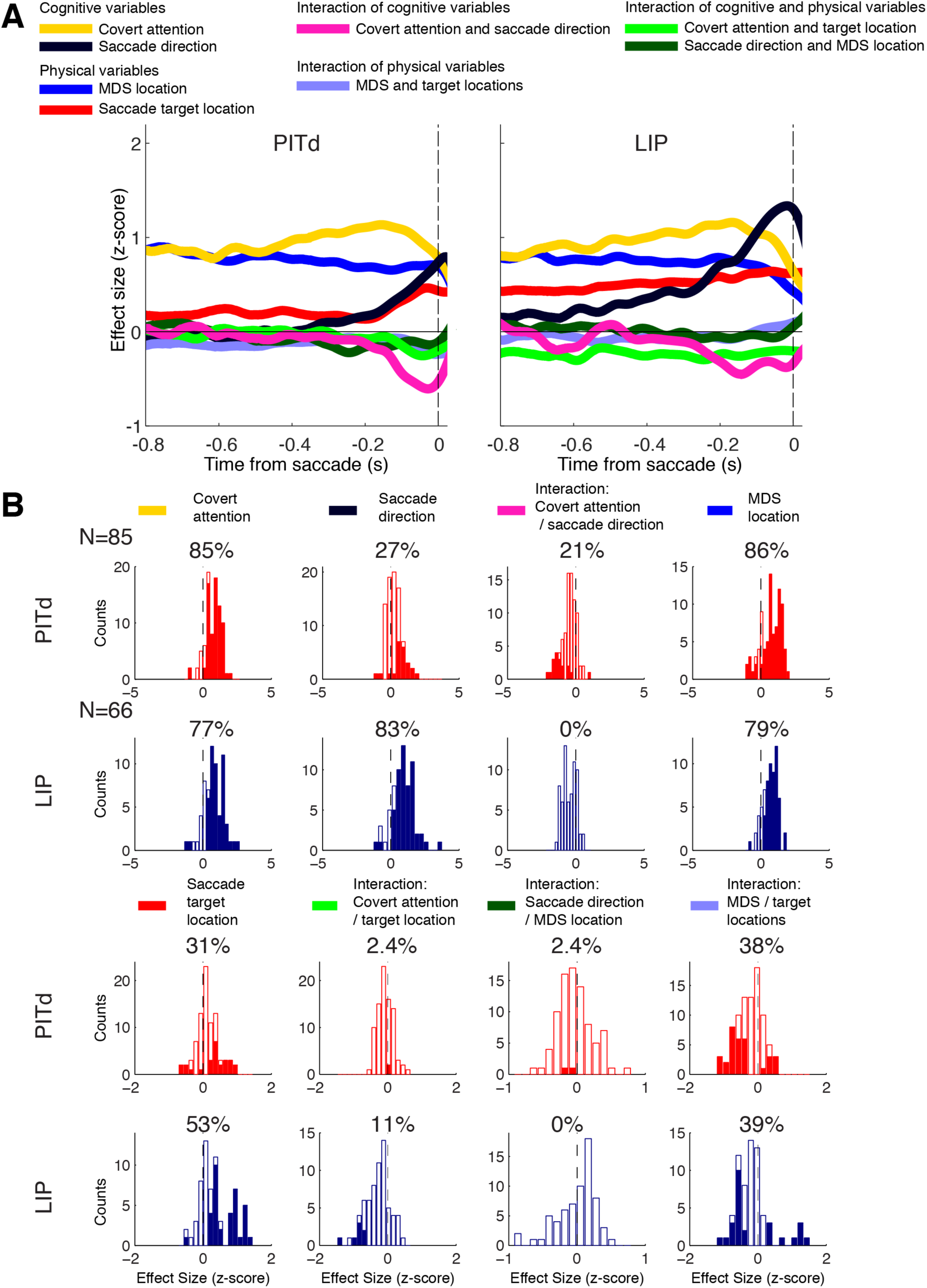
GLM Analysis (Corresponds to Figure 4B) (**A**) Mean z-scored results of GLM analysis, time aligned on saccade onset, in areas PITd and LIP. Colors indicate main effects of cognitive and physical task variables and their interactions as shown in legend. (**B**) Distribution of effect sizes (z-scored) of task variables and their interactions (see legend on top, color code from GLM time courses provided for reference) in areas PITd (red plots) and LIP (blue plots). Filled segments indicate statistically significant units (two-tailed t-test) at a false discovery rate set a q<0.05 in each area. Percent numbers on top of each plot denote fraction of statistically significant units.

**Fig S5.**
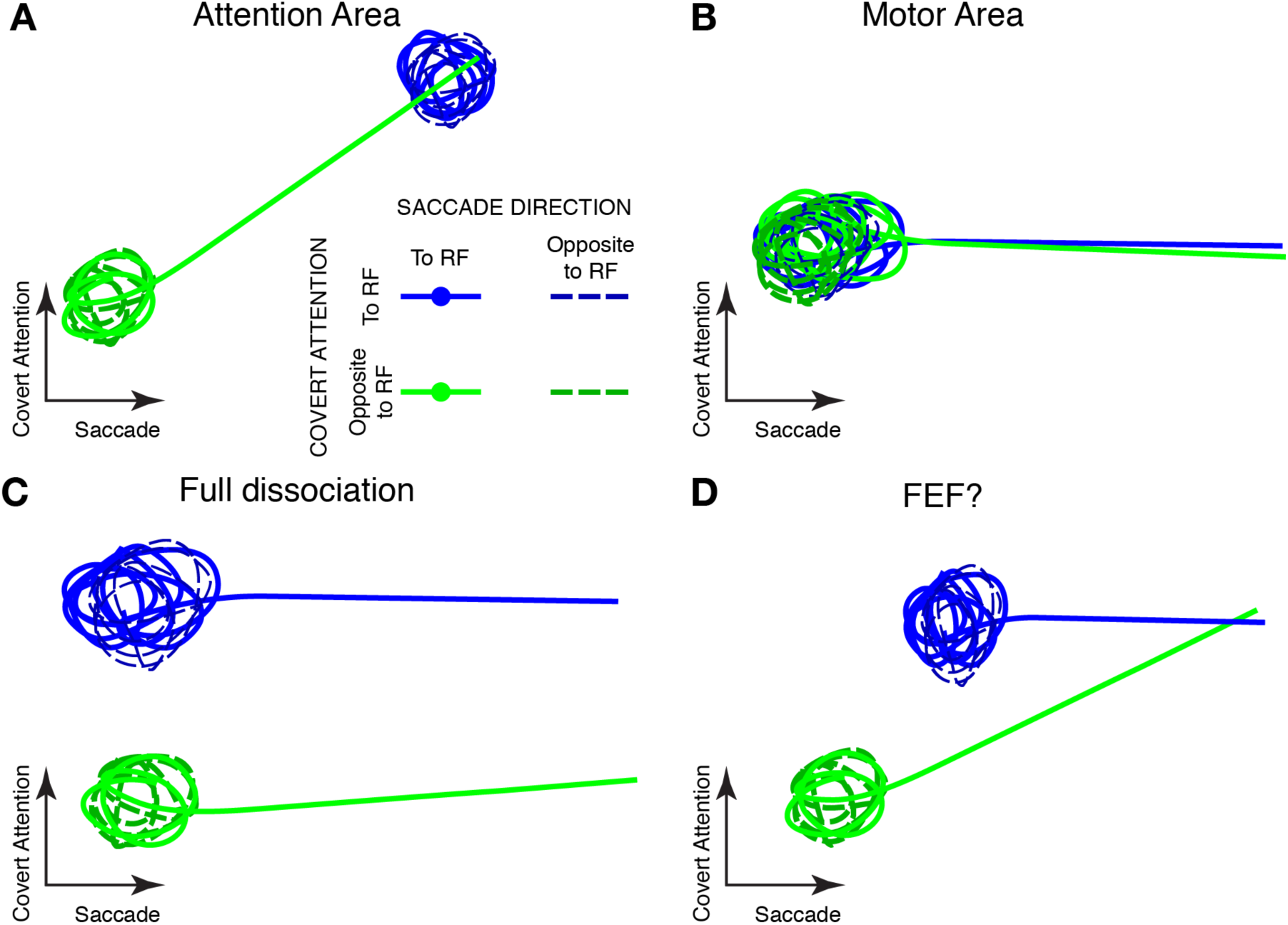
Population portraits for alternative hypothetical scenarios. (Corresponds to Figure 5B) (**A**) A population that represents only the attended location. A trace can be expected when there is reorientation of attention caused by planning a saccade to a previously unattended location. (**B**) A purely motor area, where no attention effects are seen, and saccades to the RF are always represented. (**C**) A full dissociation scenario, with independent representations for saccades and eye movements. (**D**) An area like the frontal eye fields (FEF) where a mix of purely motor and attention-motor cells has been reported (29). Conventions as in Fig. 5B.

### SUPPLEMENTAL METHODS

#### 1. Subjects and Surgical Procedures

Experiments were performed with two adult male rhesus macaques (*Maccaca mulatta*), weighing 8-10 kg. Implantation of MR-compatible headpost (Ultem; General Electric Plastics), MR-compatible ceramic screws (Thomas Recording), and acrylic cement (Grip Cement, Caulk; Dentsply International) followed standard anesthetic, aseptic, and postoperative treatment protocols.

All animal procedures complied with the US National Institutes of Health *Guide for Care and Use of Laboratory Animals* and were approved by The Rockefeller University Institutional Animal Care and Use Committee (IACUC).

#### 2. Electrophysiology experiments

##### 2.1 MRI-guided electrophysiology

Electrophysiological recordings in both animals were guided by statistical maps of the effects of covert spatial attention obtained in a previous study (20). In brief, animals were trained to perform a covert-discrimination task, related to the one used in the present study. They were required to covertly attend one out of two moving-dot surfaces (MDSs), positioned in the left and right hemifields along the horizontal meridian. Animals reported the direction (eight possibilities) of a prolonged motion event in the cued MDS, with an eye movement to one out of eight possible saccade targets. A contrast of activity on attend-left vs. attend-right conditions was used to create a statistical map which served to estimate the location of attention-sensitive hotspots in areas PITd and LIP to be targeted with recording electrodes (Fig. S1). Vertical Plastic recording chambers (Crist Instruments) were positioned vertically to reach both PITd and LIP. In monkey M2, a second lateral chamber was used in PITd recordings to avoid blood vessels. We used the Planner software (30) to calculate the angle and position of desired electrode trajectories from MR anatomical volume showing blood vessels to be avoided and functional maps. We used plastic grids, available commercially (Crist Instruments) or 3D-printed from plans created by the Planner software placed inside plastic recording chambers.

##### 2.2 Extracellular recordings

Extracellular recordings were conducted using single Tungsten electrodes (FHC, impedance 2-9 MΩ/1kHz). Electrodes were back-loaded into metal guide tubes of length set to reach, from the top of the grid holes, approximately 2mm below the dura and were slowly advanced using a manual oil hydraulic manipulator (Narishige Scientific Instruments). The electrophysiological signal was amplified and waveforms that crossed a set threshold were sorted online into separate units using multiple discrimination windows or ellipsoids defining clusters in principal component space. Spiking activity, local field potentials, eye position traces and digital triggers of task events were recorded using a Cerebus data-acquisition system (Blackrock Microsystems). Spike isolation was re-assessed offline by existence of an absolute refractory period in inter-spike interval histograms and existence of well-defined clusters in principal component space. Each well-defined cluster was treated as a ‘unit’ for the purposes of the analyses.

##### 2.3 Behavioral monitoring and stimulus presentation

Behavior was controlled and stimuli were presented using custom software ‘Visiko’ written in C++, running on a Windows PC that received and sent signals via an analog and digital input/output card PCI-DAS1002 (Measurement Computing Corporation). The software controlled juice rewards and sent triggers to the Cerebus data-acquisition system. Eye position was measured and recorded at 100Hz using an infrared eye-tracking system (ETL-200, ISCAN Inc.). Stimuli appeared on a CRT (cathode ray tube) computer monitor (36.6 x 27.4 cm; 1920 x1440 pixels; 100 Hz refresh rate) at a distance of 57 cm from the eyes.

##### 2.4 Cell selection

During recording sessions, the presence of neural activity was monitored by the sound of the amplified neural signal connected to an audio monitor and by observing the presence of waveforms on a computer screen. The consistency of transitions between gray matter, white matter and sulci as assessed from these measures was compared with expected trajectories in anatomical MR images, while expected visual, auditory or somatosensory responsiveness of areas along electrode trajectory was verified and compared with that expected from brain atlases and preceding recording sessions. In LIP recordings, activity during memory-guided saccades was used to verify that the final electrode position was correct during the first recordings in a given recording site, but this activity was not a requirement for a given cell to be recorded. In PITd recordings, the electrode was advanced until non-foveal visual responses were observed. We did not select cells based on any requirement other than visual tuning. On a typical session we would first record responses during the receptive-field (RF) mapping task, used to tune stimulus parameters (positions) in subsequent tasks. Then we recorded responses during a memory-guided saccade (MGS) task and finally the covert-discrimination task (described below). When possible, after the covert-discrimination task, we would reconfirm RF position and responses on the MGS task by recording responses a second time. We recorded single units and multi unit activity in areas PITd (85 units; M1: 25 single units (SU), 17 multi units (MU); M2: 26 SU, 20 MU) and LIP (66 units; M1: 19 SU, 19 MU; M2: 24 SU, 4 MU).

#### 3. Behavioral tasks

##### 3.1 Receptive-field mapping task

We used this task to compute a quantitative spatial map of a cell’s RF. Monkeys were required to fixate on a white central spot (diameter: 0.5 degrees of visual angle (°), fixation tolerance window: 4deg, luminance: 26cd/m^2^, CIE chromaticity: x=0272 y=0.274) on a dark gray background (luminance: 2.53cd/m^2^, CIE chromaticity: x=0.251, y=0.217). Reward was delivered after a variable period (3-3.5s) of sustained fixation. White squares (100ms duration separated by a 300ms pause, width: 2deg, luminance and chromaticity as fixation spot) were presented at random positions along an equilateral triangular lattice (spacing: 1.5deg, extent: 30deg) covering the computer screen. Experiments typically lasted 5-7min.

##### 3.2 Memory-guided saccade task

Monkeys had to make a saccade to a remembered location in the absence of a visual stimulus (Fig. 1C). Subjects initiated a trial by fixating on a central blue dot (fixation tolerance window: 5deg, diameter: 0.5deg, luminance: 6.3cd/m^2^, CIE chromaticity: x=0115 y=0.116) on a black background (luminance: 0.01cd/m^2^, CIE chromaticity: x=0.125, y=0.124). A central fixation period (300 to 600ms) was followed by a peripheral visual target presentation (duration: 200ms, diameter: 0.8deg, saccade window: 7- 10deg, luminance: 26cd/m^2^, CIE chromaticity: x=0.272 y=0.274) Target locations had a fixed eccentricity (selected to match that of the RF of the cell being recorded) and one of eight equally separated polar angles in visual space (meridians and diagonals). After extinction of the target, monkeys waited for a variable period of time (memory period, 900-1400ms) until the central fixation spot disappeared. Monkeys were rewarded with a juice drop if they made a saccade to the remembered target (∼7deg variable tolerance window) location within 400ms of disappearance of the central fixation spot. After a correct response, the saccade target briefly reappeared (100ms). Typical experiments included 100 correct trials. Measurements of the dynamics of luminance of the saccade target with a photodiode confirmed its presentation lasted 200ms, after which luminance decayed to baseline levels in a time shorter than the monitor inter-frame interval (10ms).

##### 3.3 Covert-discrimination task

Monkeys had to detect a prolonged motion event (PME) in one out of two moving-dot surfaces (MDSs) and report its direction with a saccade. Trial structure is schematized in Fig. 2A. Trials started with a 1s central fixation period (diameter: 0.25deg, fixation tolerance window: 4deg, luminance: 2.5cd/m^2^, CIE chromaticity: x=0.251 y=0.217), on a black background (luminance: 0.01cd/m^2^, CIE chromaticity: x=0.125, y=0.124). Saccade targets were present throughout the trial (annuli inner diameter: 0.2deg, outer diameter: 0.3deg, saccade window: 5-8deg, luminance: 2.5cd/m^2^, CIE chromaticity: x=0251 y=0.217). A small central rectangular cue (length: 0.25deg, width: 0.07deg, luminance and CIE chromaticity as fixation spot) appeared pointing in the direction of the future relevant MDS, staying during the whole duration of the trial. After 1s, the two MDSs (diameter: 4-5deg, mean luminance: 0.12cd/m^2^. Dot size: 0.2deg, density: 6dots/deg^2^, speed: 6deg/s, lifetime: 100ms, luminance: 1.2cd/m^2^, CIE chromaticity: x=0.245 y=0.191) appeared on opposite sides of the visual field, the cued MDS being behaviorally relevant while the other one was a distractor that had to be ignored. A fixed fraction of dots moved in a consistent direction (motion coherence 9-28%, fixed in each session). For each MDS, motion direction changed randomly and independently (24 possible directions) every 60ms for a variable number of times (1 to 60), independent for each MDS. These irrelevant translations were followed by the PME during which the direction of motion remained the same (2.7s). During the PME, dots moved into one of two opposite directions. Monkeys had to indicate the direction of motion of the cued MDS’s PME with an eye movement to the corresponding saccade target before the end of the PME. As mentioned above, motion coherence levels varied from session to session to keep performance levels around 66-75% correct. In this way we compensated for performance variability, which tended to be affected by the use of different stimulus eccentricities.

The goal of the task design was to study spatial signals related to covert attention to the MDSs and response selection with a saccade, when both are spatially separated or allowed to overlap. To do that, we had conditions in which there was only a MDS in the RF of the recorded cell, or only a saccade target in the RF, or both, or none (Fig. 2B). Each of these four stimulus arrangements included four behavioral conditions (two possible cue directions and two possible saccade directions), thus generating a total of 16 unique conditions overall. The two MDSs were always positioned on opposite sides around the central fixation point and the same was true for the two saccade targets. Depending on the trial condition, MDSs could be aligned with saccade targets or lie along a line perpendicular to them. Stimulus eccentricity was arranged to match that of the RF, polar angles were chosen between meridian and diagonal, whichever matched the RF polar angle best.

The different conditions were presented in blocks of trials that had the same stimulus arrangement and cued MDS, but randomized PME timing and direction. A block was completed when six trials had been performed correctly.

Several aspects of task design ensured that monkeys needed to gather relevant information from the cued MDS as opposed to randomly guessing, getting useful information from the distractor MDS, or responding in an impulsive or stereotyped way. First, in order for responses to be considered correct monkeys had to make a saccade only after the PME on the cued MDS started, which reduced chance performance levels well below 50% if an eye movement was made at random times (See Fig. 2C). Second, the onset time of the PME varied within a long time window, which made it difficult for monkeys to randomly guess when a response was appropriate. Third, 10% of trials were “catch” trials that occurred randomly within the blocks, where no prolonged motion event occurred in the cued MDS, and monkeys were rewarded for maintaining fixation until the end of the trial. Fourth, the PMEs in the cued and distractor MDSs were uncorrelated both in time and motion direction, ensuring no useful information about either timing or motion direction could be extracted from the distractor MDS. Fifth, during training sessions preceding recordings only, incorrect trials were repeated until the monkey provided a correct response so as to make stereotyped responses an inviable strategy. In recording sessions, incorrect trials were repeated in non-consecutive trials instead. Sixth, trials with incorrect responses were punished with an increased inter-trial time. Seventh, reward magnitude was incremental, i.e. increased after a successful trial and reset to initial levels after a mistake, encouraging monkeys to correctly perform many trials in a row.

#### 4. Analysis of behavior during the covert-discrimination task

##### 4.1 Analysis of overall performance

To study whether behavioral responses were reliably paired with events in the cued and/or distractor MDSs in individual trials, we categorized the responses in each trial as being consistent with events in the cued MDS, the distractor MDS, both of them, or none. A response was consistent with events in a MDS if and only if the saccade occurred within 2.7s of the time of PME onset (tPME_OT) of that MDS and the saccade direction matched that of the PME. Trials consistent with the cued MDS or both MDSs corresponded to correct trials, as opposed to trials consistent only with the distractor MDS or none of them. Only trials in which monkeys made a response to a saccade target were considered in this and subsequent behavioral analyses; trials aborted by breaking central fixation or by making a saccade to one of the targets before MDS onset were excluded. For every experimental session, we compared the observed proportion of responses of each type with those from a null distribution generated by randomly permuting the trial number of the PMEs to witch behavioral responses were paired (permutation test). For each response type, the fraction of permutations that had a percent of responses of that type more extreme than the one observed determined the statistical significance. We used 10^5^ random permutations for the computation of statistical significance.

##### 4.2 Analysis of detection behavior

Our measure of detection performance was the percentage of successfully detected PMEs of the cued or distractor MDS, i.e., the percentage of trials whose response time (RT) was between 0 and 2.7s after the prolonged motion event onset time (tPME_OT). It is important to realize that even a high percentage of successfully detected PMEs can be achieved if the monkey learns the temporal statistics of tPME_OT and waits until the mean or median tPME_OT has passed before making a saccade. This is also bound to be the case if, for instance, monkeys respond to the cued MDS’s PME in each trial with a considerable delay, in which case they will tend to respond after the distractor MDS tPME_OT on many trials too. Because of these considerations, we looked for evidence that, in each trial, the monkeys responded to the specific PME of that trial rather than the overall tPME_OT statistics. Since the cued and distractor MDSs have a PME each, both are studied separately. The significance of the detection behavior was estimated using a permutation test, where the percentage of successfully detected PMEs was compared to the distribution obtained by randomly permuting the tPME_OT of all trials. The fraction of permutations that resulted in an increase of detection behavior relative to the one actually observed provided the statistical significance of the discrimination behavior. We used 10^5^ random permutations for each computation of statistical significance.

##### 4.3 Analysis of discrimination behavior

Our measure of discrimination behavior was the percentage of successfully detected PMEs that were also successfully discriminated. In all trials, there were two saccade targets, each with 50% probability of being correct. A binomial test was used to compute the probability that a discrimination behavior better than the one observed could arise by chance.

##### 4.4 Analysis of correlation between timing of trial events and behavioral response

To study if monkeys somehow integrated the PME of cued and distractor MDSs in their detection behavior, we looked at the correlation of the RTs measured relative tPME_OT of the cue and distractor MDSs. Since both quantities share a common trigger, a positive correlation between them is to be expected. The hypothesis to test, then, is that the observed correlation is higher (monkeys combining events from both MDSs) than the one simply produced by triggering the tPME_OT on a dummy event that shares the overall distribution of the RTs. We generated dummy triggers using RTs from randomized trials and computed a distribution of correlation values (permutation test). The statistical significance of the observed correlation value was the fraction of correlation values under the null hypothesis that were lower than the one observed. We used 10^5^ random permutations for the computation of statistical significance.

#### 5. Analysis of neural responses

##### 5.1 Receptive-field mapping

For each stimulus position, the mean firing rate across repeated presentations (including only those in which there was successful central fixation) was calculated using a temporal window from 50 to 150ms after stimulus onset. Firing rates were interpolated to all positions in visual space via a radial basis method. Results were smoothed via convolution with a Gaussian kernel (2deg FWHM, truncated at 3deg). The RF center was estimated in the following way. First, the smoothed map was converted to z-scores. Then, points with z<0 where removed. The largest connected component of the resulting map was computed. Its center of mass was taken to be the RF center.

##### 5.2 Memory-guided saccades

All analyses were performed in coordinates relative to the RF, either estimated from the RF mapping task or, in rare exceptions when the acquired map was deemed too noisy for an accurate estimate of the RF center, from the target position that elicited the largest visual response. For each trial, we calculated a spike-density function (SDF) by convolving each spike (considered as a Dirac delta function) with a Gaussian kernel (σ=25ms, truncated at 100ms). Activity was aligned separately to two events: target onset and saccade onset. Target-onset-aligned activity included time from 500ms before target onset until 500ms before saccade onset. Saccade onsets were defined as the time the eye position left the fixation window (width: 2 degree). Saccade-onset-aligned activity included spikes from 400ms after target onset until 200ms after saccade onset. Because trials have different durations, a variable number of trials contribute to the mean time course in a given task condition, but similar across task conditions for a given time point. For quantifying the strength of effects across the population, the activity of each cell was z-scored as follows. First, for each of the 8 target positions, the mean time course across repeated trials was computed, to ensure all conditions were equally represented independently of the number of trials available. Second, the mean activity and standard deviation across all 8 target positions and time points was computed.

Third, the mean was subtracted from the SDF, which was then divided by the standard deviation. To compare the distribution of tuning strength during the memory period, we used a time window from -600ms to -400ms relative to saccade onset in trials in which the target was positioned inside the RF and trials in which it was opposite to it. We quantified the strength of tuning of activity to the target position by fitting a generalized linear model (GLM) with a Poisson noise distribution with target presence in the RF as the only predictor. We used two-tailed t-tests to quantify the significance of results as implemented in MATLAB’s GLM-fitting function *glmfit*. To estimate the percentage of units in each brain area with significant tuning, while controlling for comparisons across multiple units, we set the false discovery rate (FDR) to q<0.05 using the two-stage Benjamini-Krieger-Yekutieli procedure (31). Since results from both monkeys were consistent, we pooled together units from both for analysis.

##### 5.3 Covert-discrimination task: general procedures

As above, all analyses were done in coordinates relative to the RF center determined above. SDFs were calculated as in the MGS task. Activity was aligned separately to two events: MDS onset and saccade onset, the latter corresponding to the time point where gaze leaves the fixation window. The MDS-onset-aligned analysis window ranged from 1s before MDS onset to 500ms before saccade onset. Saccade-onset-aligned activity included spikes from 300ms after MDS onset until 600ms after saccade onset. As in the MGS task analyses, activity of each unit was z-scored for estimating effects across the population: average time courses were calculated for each of the unique 16 conditions, global mean and standard deviation were calculated across all time points and conditions. Then, SDF in each trial and time point had the global mean subtracted and was divided by the global standard deviation.

As above, since results from both monkeys were consistent, we pooled together units from both for analysis.

##### 5.4 Covert-discrimination task: generalized linear model

To quantify the effect of task variables on the responses of each recorded unit, we used a multi-variable, generalized linear model (GLM) with a Poisson noise distribution. Activity was described as

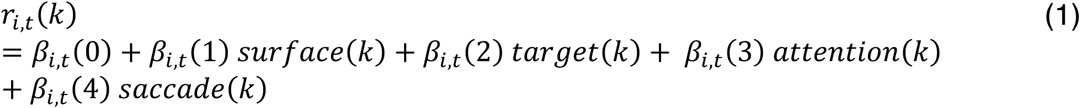

where 𝑟_!,#_(𝑘) is the firing rate of neuron *i* at time *t* in trial *k*, 𝑠𝑢𝑟𝑓𝑎𝑐𝑒(𝑘) indicates the presence of a MDS in the RF (+1/2: present, -1/2: absent), 𝑡𝑎𝑟𝑔𝑒𝑡(𝑘) refers to the presence of a saccade target in the RF (+1/2: present, -1/2: absent), 𝑎𝑡𝑡𝑒𝑛𝑡𝑖𝑜𝑛(𝑘) denotes if covert attention is directed to the RF (+1/2: to RF, -1/2: opposite to RF, 0:

MDS not present in RF), 𝑠𝑎𝑐𝑐𝑎𝑑𝑒(𝑘) specifies saccade direction (+1/2: to RF, -1/2: opposite to RF, 0: saccade target not present in RF).

The regression coefficients 𝛽_!,#_(𝜈), for 𝜈 = 1, … ,4 quantify how much a given task variable (𝜈 = 1: MDS presence, 𝜈 = 2: target presence, 𝜈 = 3: covert attention, 𝜈 = 4: saccade direction) contributes to the observed firing rate. The first regression coefficient (𝜈 = 0) reflects the time-dependent component of the firing rate that is not dependent on any of the task variables. Equation (1) reflects the simplest regression of the firing rate, but we critically also included additional quadratic terms between experimental variables quantifying deviations from the simple sum assumed in (1) that could be explained by interactions between predictors. We also included nuisance terms (not shown here for simplicity) that captured potential effects of covert attention to different positions when the MDS was not inside the RF and an analogous term for saccade direction when saccade targets were not positioned in the RF. All beta values obtained were converted from spikes/second to z-scores for each unit via normalization by the standard deviation described above (section 5.2).

The GLM was computed independently at each time point to display the dynamics of the different effects. For quantification the distribution of effects across cells, we performed the regression using activity collapsed in fixed time windows: 200 to 0ms before saccade onset for saccade tuning and its interactions, and 800 to 500ms for all other effects and associated interactions. All statistical tests were two-tailed t-tests as implemented in MATLAB’s GLM-fitting function *glmfit*. To estimate the percentage of units in each brain area where each of the effects was significant, while controlling for comparisons across multiple units, we set the false discovery rate (FDR) to q<0.05 using the two-stage Benjamini-Krieger-Yekutieli procedure (31).

##### 5.5 Covert-discrimination task: Decoding analyses

We quantified the presence of distributed information in the population about covert attention and saccades using a decoding approach, in which we trained a decoder using the activity of pseudo-populations of units from each area. We used linear support vector machines (SVMs)(32) to classify task conditions, which, while simple, can be interpreted as performing computations of a downstream neuron and are particularly suitable for multi-dimensional representations (23). In that sense, it is said that linear SVMs are sensitive to information that is explicitly represented in the population. Decoding results obtained this way provide a lower bound for the amount of information available at the population level, which can in principle be extracted using more complex algorithms.

Decoding analyses were performed using the Neural Decoding Toolbox (33) (version 1.0.2), a MATLAB package implementing neural population decoding methods. Activity of all units was considered relative to their receptive field, so as to form a collection of units that encode information about the same portion of space, similar to what is expected from cells belonging to the same cortical column. We performed these analyses on data aligned to saccade onset, binned every 100ms. We restricted decoding to units for which at least 6 repetitions from each task condition used in each analysis was available. Each subpopulation was composed of the same number of units, randomly sampled with replacement from the pool of units available from each area. We followed a cross-validation approach, in which a subset of the data was used to train the decoder, while the remaining trials were used to test the decoding accuracy, as a measure of the reliability of the learned relationship between neural activity and experimental condition.

For each unit, data were randomly selected from 6 trials from each of the relevant conditions. For each of these trials, data from 20 units was concatenated to create pseudo-population response vectors (that is, data from units recorded mostly on separate sessions but treated as if they had been recorded simultaneously). The pseudo-population vectors were grouped into 6 splits of the data, each containing a response vector for each relevant condition. A linear SVM was trained using 5 splits of the data and the performance of the classifier was tested using the remaining split of the data. Before providing the data to the classifier, data from each unit was z-scored (using parameters estimated from the training set), to prevent neurons with high firing rates from dominating the representation of information. The same procedure was repeated 6 times, each time selecting a different split of the data as test set. In other words, a 6- fold leave-one-split-out cross-validation procedure was used. As a measure of performance, we calculated the fraction of test examples that were correctly classified (i.e., zero-one loss measure). To get a more precise estimate of the decoding performance as well as an estimation of the variability of the results that would arise as a consequence of selecting different subsets of neurons drawn from the same original population, we repeated the decoding procedure 200 times, using a bootstrap resampling method. On each repetition, units were drawn randomly from the available pool with the possibility of drawing the same unit more than once (random sampling with replacement). The results we report are averaged across the 200 repetitions.

To determine statistical significance of results, we used permutation tests. Decoding procedures were repeated as before, but labels of experimental condition labels were randomly permuted before supplying them to the decoder. We used 50 rather than 200 repetitions for each permutation to reduce computation time. This makes the test more conservative since it overestimates the variance of the null distribution, relative to the use of 200 repetitions. As a consequence, the probability of a type II error was increased, but not that of a type I error. p-values were adjusted for comparisons across multiple time points using a Holm-Bonferroni procedure to control the family-wise error rate (34).

##### 5.6 Covert-discrimination: Extraction of task relevant dimensions

To visualize the dynamics of the population along dimensions most informative about task variables, we used a recently developed dimensionality reduction technique, targeted dimensionality reduction(24), which proceeds as follows. The average SDFs from all cells in a given area are concatenated to form a population SDF vector. SDF vectors are “de-noised” by performing a principal component analysis (PCA) and reducing the dimension of the vector to a smaller number of informative dimensions. In our case we used 12 principal components (PCs), following Mante and colleagues(24). Targeted dimensionality reduction isolates a dimension associated with any task variable of interest by considering regression coefficients like those of equation 1 (section 5.4) of a collection of units as a vector. The regression coefficients of all units for the task variable of interest are concatenated forming a vector, the dimension of which equals the number of units. This vector is then projected to the same PC space computed above from SDFs. Since regression coefficients evolve in time, so does their associated vector. The direction associated with a task variable of interest is the direction of its regression coefficient vector in PC space at the time point where the norm of the vector reaches its maximum value. It should be noted that when more than one variable of interest is isolated, the associated directions are in general not orthogonal. This is problematic because the projection of activity to the subspace spanned by these directions, although unique, is not simply the independent projection to each of the computed directions. This can be solved by an orthogonalization procedure (such as a Gram–Schmidt process), which leaves the spanned subspace unchanged. The multidimensional SDFs can then be linearly projected into the resulting axes for visualization.

We made a minor but important modification to the targeted dimensionality reduction technique. The priority map hypothesis proposes that neural activity in the presence of multiple task factors is roughly the sum of responses to each task factor in isolation. To test this hypothesis, regression coefficients from trials with MDSs *alone* inside the neurons’ RF were used to compute a direction associated with covert attention. Analogously, regression coefficients from trials with saccade targets *alone* in the RF were used to isolate a direction associated with saccade direction. We then used these directions to project the population SDFs when *both* MDSs and saccade targets were present in the RF. This modification has the dual advantage of testing the proposed encoding scheme for priority maps and, in addition, serving as a cross-validation procedure ensuring the selected directions do not reflect the particular realization of the noise in the conditions plotted. All calculations were done using neural activity temporally aligned to saccade onset.

### BEHAVIOR RESULTS

We determined whether monkeys preformed the task using the central cue to select the cued MDS and ignored the distractor MDS. Trials explicitly aborted by breaking central fixation with an eye movement to none of the saccade targets were disregarded in behavioral analyses presented here. There were 16471 (M1) and 17769 (M2) trials in which monkeys provided a valid response from the 41 (M1) and 45 (M2) experimental sessions where neural activity was recorded, resulting in about 400 valid trials per session. To receive reward, subjects were required to respond within 2.7s after the onset of the prolonged motion event (PME) of the cued MDS (successful detection), and to generate a saccade in the direction of the PME (successful discrimination). We therefore subdivided our analysis of behavior in two parts beyond overall performance: detection and discrimination. Detection was considered successful when a saccade occurred within 0 and 2.7s after PME onset to one of the saccade targets; discrimination, when, in addition, the correct saccade direction was selected. Discrimination is thus defined for trials with successful detection only. Note that, for analysis purposes, both detection and discrimination can be defined relative to either the cued MDS or the distractor. In this sense, a successful trial is one where both detection and discrimination were successful, and the overall performance is the percentage of successful trials out of the ones included in the analysis.

We first studied overall performance (joint detection and discrimination of PME) by considering the percentage of responses that were consistent with the events in either the cued MDS, both MDSs, the distractor, or none (Fig. S2A, see Methods). Note that the first two cases correspond to correct trials and the last two to error trials. For both monkeys in every experimental session, percentage of trials consistent with the cued or both MDSs were above chance levels (permutation test: p<0.05 in every session, Holm-Bonferroni corrected for number of sessions considered) and trials consistent with only the distractor MDS or none were below chance levels (permutation test: p<0.05, Holm-Bonferroni corrected for number of sessions, except for one session in monkey M2 with distractor-consistent trials not distinguishable from chance). These observations suggest monkeys successfully used selected information from the cued MDS and ignored the distractor.

We then asked if the same difference between cued and distractor MDSs existed in the separate detection and discrimination aspects of the task. We first analyzed detection, which refers specifically to the timing of responses relative to the onset time of PMEs (tPME_OT). We measured the probability that successful detection would be improved if the tPME_OT of all trials were randomly permuted. For discrimination, since both PME directions are equally likely, chance performance followed a binomial distribution. Both detection and discrimination behaviors with respect to the cued MDS differed significantly from chance levels in every experimental session (p<<0.05, Holm-Bonferroni-corrected for the number of sessions; see Methods), but with respect to the distractor MDS they did not differ significantly from chance levels in any session (p>0.05, Holm-Bonferroni corrected for the number of sessions; see Methods). This is illustrated in Fig. S2B, which shows the detection and discrimination performance, averaged across experimental sessions, of both monkeys with respect to the cued and the distractor MDSs. Together, the results strongly suggest that both monkeys successfully used the cue to guide their behavior and ignored the distractor MDS in terms of overall performance as well as regarding separate detection and discrimination.

We further considered the possibility that occurrence of PME in the distractor surface MDS could help detection of a simultaneously occurring PME in the cued surface. For this purpose we studied the distribution of response times (RTs) relative to cued and distractor tPME_OT and their correlation. tPME_OT values ranged from 0 to 3.6s relative to the onset of the MDSs and were independent for cued and distractor MDSs. Figure S2C shows the frequency distributions of tPME_OT values, separately for cued and distractor MDSs (left and right panels, respectively). tPME_OT values were chosen from a uniform distribution. Yet the empirical distribution for the cued-MDS tPME_OT is slightly biased towards late times. This is because stimulus conditions of error trials were repeated at later times in order to prevent monkeys from systematically avoiding completion of certain conditions (see Methods). Since monkeys tended to perform worse on trials with late cued-MDS tPME_OT values, these are overrepresented in the empirical distribution. Figure S2D shows the distributions of RTs in monkeys M1 and M2. RTs varied widely, for both animals, over several seconds. Figure S2E illustrates the statistical relationship between RTs and the tPME_OT of both cued and distractor MDS in monkeys M1 and M2. Figure S2E (left panels) show the joint distribution of RTs and tPME_OT of the cued MDS is highly structured, with most RTs occurring systematically after their associated tPME_OT. This is in stark contrast with Fig. S2E (right panels), which shows the joint distribution of RTs and distractor tPME_OT is appreciably less structured and close to homogeneous. It is conceivable however, that information regarding the distractor tPME_OT and the cued MDS tPME_OT are combined to facilitate detection. Figure S2F (left panels) shows the joint distribution of RTs measured relative to the cued and distractor tPME_OT. The majority of response events occur within a tight time range relative to cued MDS, observable as a horizontal stripe in the joint distribution plot. A positive correlation is to be expected if only for the fact that both quantities share a common trigger, which is shown in Fig. S2F (right panels) where RTs from random trials are used to compute the joint distribution rather than same-trial RTs. We verified the correlation between RTs relative to cued and distractor MDS was not higher but actually significantly lower than the value obtained when RTs are randomized (M1: r=0.26, p<<0.05 M2: r=0.33, p<<0.05, permutation test, see Methods). Hence, successful performance decouples responses relative to each surface.

Together, the results strongly suggest that both monkeys successfully used the cue to guide their behavior and ignored the distractor MDS in terms of overall performance as well as regarding separate detection and discrimination.

